# Serotonin signaling in the enteric nervous system and connection to autism spectrum disorder: a translational mathematical model

**DOI:** 10.1101/2022.10.11.511819

**Authors:** Irina Kareva

## Abstract

While the causes of autism spectrum disorder (ASD) remain unclear, some studies have shown that serotonin-mediated effects on enteric nervous system (ENT) correlate with ASD-like behavioral phenotype in mice. Introduced here is a mathematical model of interactions between gut serotonin and its impact on ENT. The model was used to identify three key factors that affect ENT size, namely, serotonin production, its clearance, and its ability to act as a growth factor on ENT. The model was used to reproduce experimentally reported results from a mouse model by Margolis et al. (2016), which connected serotonin-mediated ENT hypoplasia to an ASD phenotype. The proposed mathematical model was used to scale the quantified relationship from mice to humans to show how the combination of these three factors can translate to a quantifiable metric that could potentially be correlated to ASD spectrum. A detailed discussion of how ENT hypoplasia could mechanistically affect CNS activity concludes this paper.

## Introduction

Autism spectrum disorder (ASD) is a complex heterogeneous neurodevelopmental condition characterized by varying degrees of challenges in social communication and interactions, often coupled with repetitive behaviors, as well as impaired ability to learn and behave (1,2). While causes for ASD remain unknown, several twin studies suggest high degree of genetic heritability, with correlations among monozygotic twins being significantly higher than those for dizygotic twins on multiple ASD measures (3,4). Furthermore, ASD is often associated with high prevalence of gastrointestinal (GI) issues (5), and it has been suggested that serotonin (5-hydroxytryptamine, or 5-HT) may be a factor that is affecting both sides of this equation (6). In fact, elevated serotonin levels in the blood were one of the first biomarkers correlated with autism, with a quarter to a third of individuals with ASD exhibiting hyperserotonemia (7,8). Over 95% of serotonin is synthesized in the gut (9); blood serotonin is found in platelets (10), which do not synthesize it but utilize a 5-HT transporter SERT to take it up as they circulate through the gut (11,12). Hyperactivity of SERT, which is encoded by a variant of Slc6a4 gene, has been associated with both ASD and interestingly also with obsessive-compulsive disorder (OCD).

A possible mechanistic link between serotonin specifically in the gut and ASD has been studied in an intriguing experiment by Margolis et al. (13). In the gut, serotonin functions not only as a neurotransmitter but also as a growth factor (14,15) by promoting neurogenesis of the enteric (gut) nervous system (ENT). In (13), the authors evaluated the hypothesis that SERT hyperactivity in a mouse model can drive structural and functional GI deficits due to altered serotonin signaling in the gut. For that, the authors compared the impact of SERT signaling on ENT size for mice with hyperactive SERT (Ala56 mice), who exhibit whole blood hyperserotonemia as well as brain and behavioral abnormalities, with wild type mice, and with SERT receptor knock out mice (SERTKO). The authors show that due to increased activity of SERT in Ala56 mice, the size of the ENT can be up to 60% smaller at 6–8 week time point compared to WT mice, since increased clearance of serotonin from the gut due to SERT hyperactivity prevents serotonin from acting as a growth factor on ENT. These animals tended to exhibit repetitive behaviors, delayed communication, and deficits in social interactions. Furthermore, the authors showed that the deficits could be rescued by administering highly selective 5-HT4 agonist, prucalopride, during critical periods of neurodevelopment, which normalized enteric neuronal numbers in the SERT Ala56 mice and alleviated some of the social deficits.

The goal of this work is I want to explore more deeply the possible factors that connect gut concentration of serotonin, SERT expression and ENT size. This is achieved through a mathematical model aimed to reproduce key features of the experiment conducted by (13). The model is used to identify three key factors that can affect the relationship between serotonin and ENT. A possible scaling of these results from mice to humans is proposed, followed by simulations that take into account changes in serotonin production that have been observed in neurotypical children compared to children with ASD. A discussion of possible connections between hypoplastic ENT and its impact on the brain, as well as possible implications of this hypothesis, concludes this work.

## Methods

For the purposes of reproducing qualitatively the relationships described in the experiment by (Margolis et al. 2016), consider a 2-dimensional model that describes the dynamics of serotonin in the gut Sg(t) and its impact on development of the enteric nervous system N(t). Assume that there exists a normal gut serotonin synthesis rate S_0_, and that Sg is cleared from the gut at a SERT-receptor-independent rate sout; clearance rate can also be increased as a function of SERT receptor and is accounted for by parameter sert.

The enteric nervous system N(t) is assumed to increase proportionally to Sg that still remains in the gut proportionally to the amount of available serotonin per volume of N according to saturating function 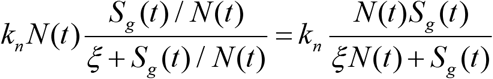, where parameter k_n_ is represents the rate of conversion of gut serotonin into ENT volume, capturing the overall effect of gut serotonin as a growth factor, as indicated by experiments (Margolis et al. 2016). Assume also that ENT can become pruned over time at some rate *n_out_*, since it cannot grow indefinitely.

The resulting system of equations becomes

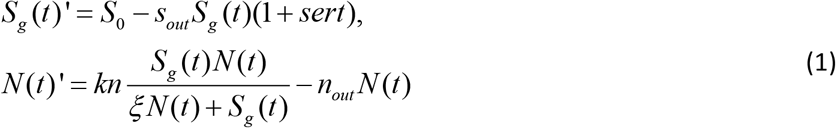

Parameters used in simulations are summarized in Table A1. Values were chosen to qualitatively capture the experimentally observed behavior and do not necessarily represent physiological values.

**Table 1.** Parameter values used in simulations for both mouse and human dynamics.

| Parameter | Description | Value (mouse) | Value (human) |
| --- | --- | --- | --- |
| $k_n$ | Rate of conversion of serotonin into ENT | 0.5 | 10.1 |
| $n_{out}$ | Rate of normal ENT pruning (to have the system reach equilibrium) | 0.1 | .99 |
| $S_0$ | Serotonin synthesis rate in the gut | 0.2 | 1 |
| $k_{deg}$ | Gut serotonin degradation rate | 1.386 | 1.386 |
| $k_{syn}$ | $S_0 * k_{deg}$ | 0.2772 | 1.386 |
| $sert$ | Rate of sert-mediated serotonin uptake | 0 if knockout (KO)<br>0.24 if normal<br>1.1 if sert+ | 0.4 for norm |
| $\xi$ | Half-maximal concentration of serotonin needed for gut serotonin-mediated ENT growth | .19 | 0.9 |
| $S_{out}$ | Serotonin clearance from the gut not through sert | 0.0035 | 0.01 |

## Results

### Reproducing the qualitative relationship between gut serotonin, ENT size and serotonin clearance in mice

Firstly, we want to qualitatively reproduce the reported relationship between serotonin in the gut, SERT receptor expression and relative size of ENT at approximately t = 50 d (mice in (13) were examined at 6-8 weeks). In Figure 1, the simulation was run for 150 days, depicting the 3 cases for SERT expression: KO (value of parameter *sert* = 0), WT (*sert* = 0.24) and Ala56, also notated as SERT+ (*sert* = 1.1). Values of parameter *sert* were chosen in such a way as to capture 100% ENT as the norm (WT), 120% ENT relative to norm for KO, and 60% ENT relative to the norm for SERT+ at t = 50 d as reported in Figure 1 (A,E) and Figure 2 (A,E) in (13).

**Figure 1.**
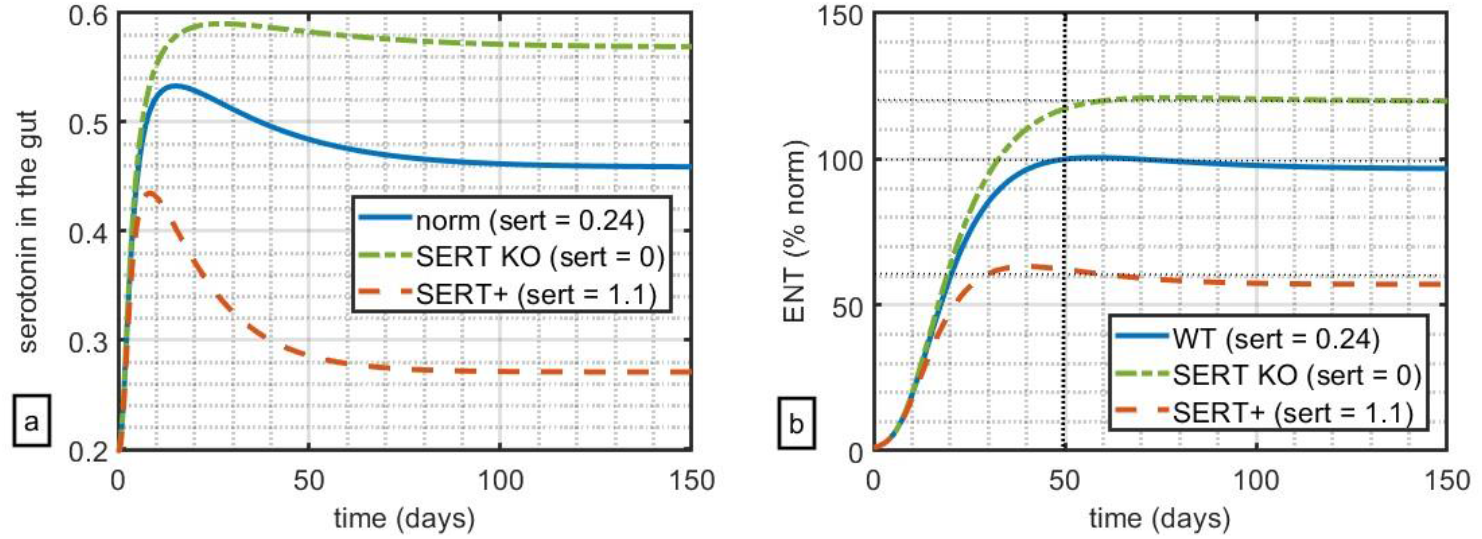
Impact of relative levels of SERT expression on (a) simulated serotonin in the gut and (b) size of ENT relative to WT. At t=50 days, WT mice had 100% ENT, with SERT KO mice at 120% of ENT, and SERT+ mice at 60% ENT relative to WT.

**Figure 2.**
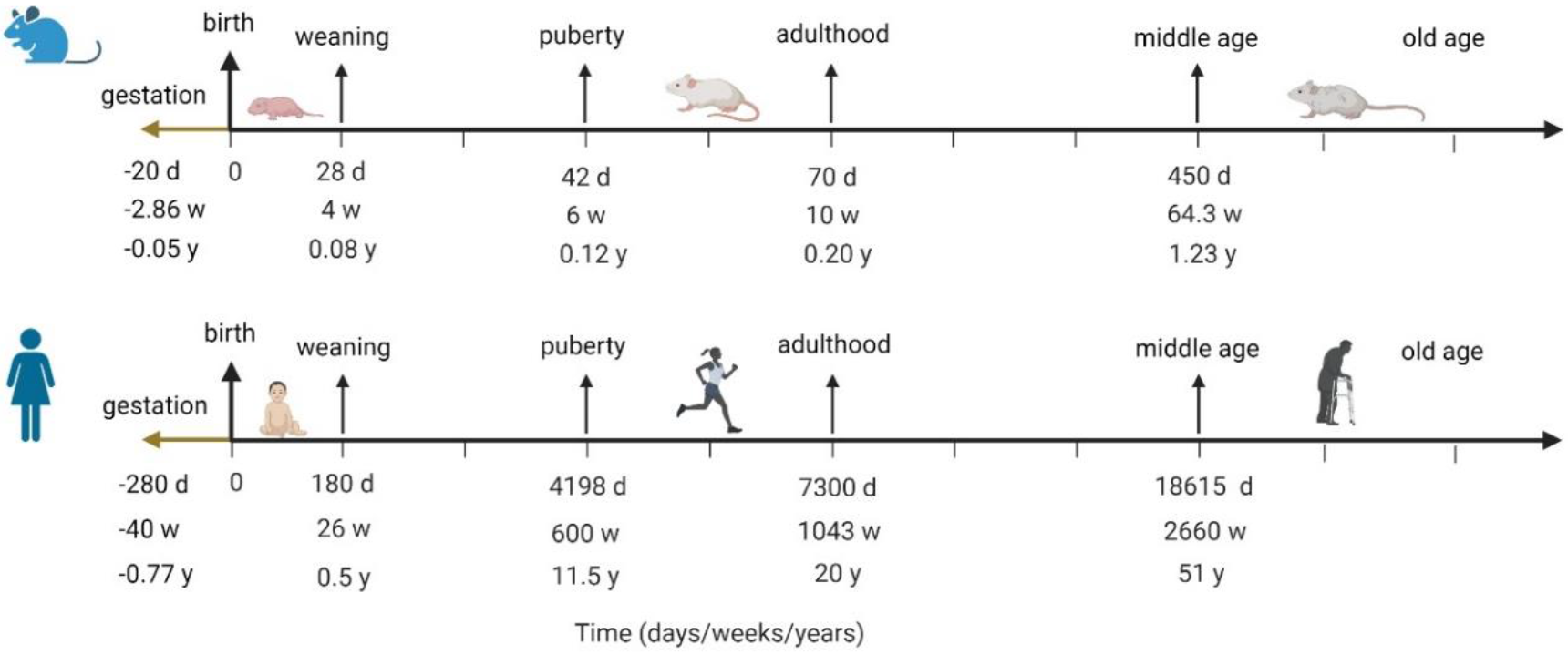
Aligning human and mouse lifespans by developmental milestones; the figure visualizes analysis done by Dutta et al. in (16).

These simulations suggest that in this model, just variation of parameter of SERT-mediated clearance can qualitatively capture the key feature of serotonin-ENT-SERT relationship reported in (13).

### Translating model dynamics from mouse to human

Of course, the question of inter-species translatability is always critical for extrapolating any insights from animal models into humans. Before this question can be addressed, it’s important to understand how mouse lifespan maps onto the human lifespan, since ASD is typically diagnosed in children before 5 years old. Instead of assuming that each mouse day has a uniform conversion to a human day throughout the animal’s lifespan, it is more instructive to align the two lifespans by key developmental milestones, from gestation, to weaning, to puberty, to adulthood and into old age, as has been done by Dutta et al. (16). Figure 2 visualizes the analysis done in (16), aligning mouse and human lifespans by developmental milestones.

Margolis et al. (13) report data that was collected in mice at 6-8 weeks, which, according to the mapping in Figure 2, corresponds to humans of about 11-13 years old. It is unknown if these animals exhibited ASD-like behaviors around week 5, which would be closer to human age of around 5 years old, by which time we would already observe ASD-like behavior, although it is possible. If these relationships indeed translate, and if the relative size of ENT is indeed correlated with ASD-like phenotype, then the differences in ENT would be observed by age 5. The model for projected human analysis is therefore re-parametrized to capture similar relationships between ENT size for non-ASD (parametrized to 100%) vs ASD kids by t = 5 years, with the understanding that it is possible for ASD symptoms to manifest much earlier.

Furthermore, interestingly, Chugani et al. (17) reported difference in serotonin synthesis capacity in ASD vs non-ASD kids, showing that for non-autistic children serotonin synthesis capacity increased more sharply and to higher values in early life but declined towards adult values after 5 years old, while in autistic children, it increased more gradually until 15 years old but did not decline as much as it did for non-ASD children. Within the frameworks of the proposed model, these differences can be captured as a term for serotonin production, with the assumption that at least the pattern of relative differences in serotonin synthesis capacity reported in Chugani et al. (17) mirrors that of serotonin synthesis in the gut, a hypothesis that requires further experimental validation. A mathematical expression to describe change in serotonin synthesis for ASD and non-ASD children is described in Appendix B. The projected serotonin dynamics over time for ASD and non-ASD children using this function is shown in Figure 3.

**Figure 3.**
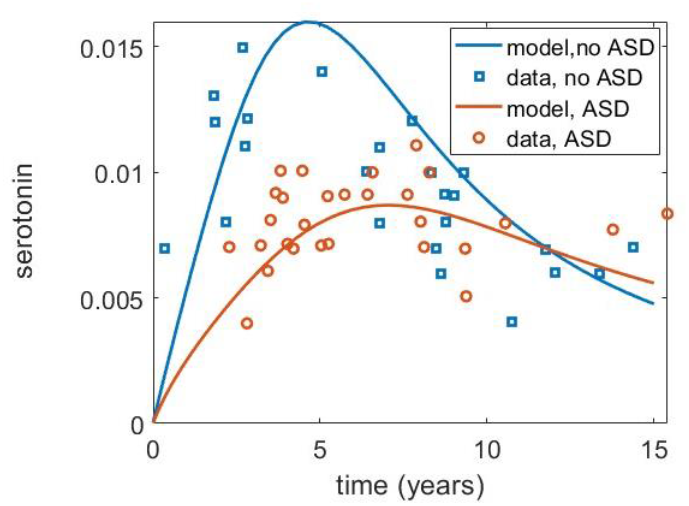
Differences in serotonin synthesis capacity over time for ASD and non-ASD children as reported by Chugani et al. (17) and captured by the model.

### Impact of individual mechanisms on ENT

It can be seen from the model, and predicted intuitively, that in addition to serotonin production (here captured by parameter *a* in Equations (A1) and (A2) in Appendix, there are two other key components that could qualitatively affect ENT size, namely, serotonin clearance through activity of SERT, drugs, or any other receptor that is involved in clearance of serotonin from the gut, preventing it from acting as a growth factor (here captured as parameter *sert*); and the rate of ENT growth as a function of serotonin (here captured by parameter k_n_ in Equation (A1) in Appendix). In Figure 4, all the three factors are varied to assess their relative impact on ENT size relative to norm (blue, solid) by time = 5 years, including the dynamic human serotonin production function from Figure 3.

**Figure 4.**
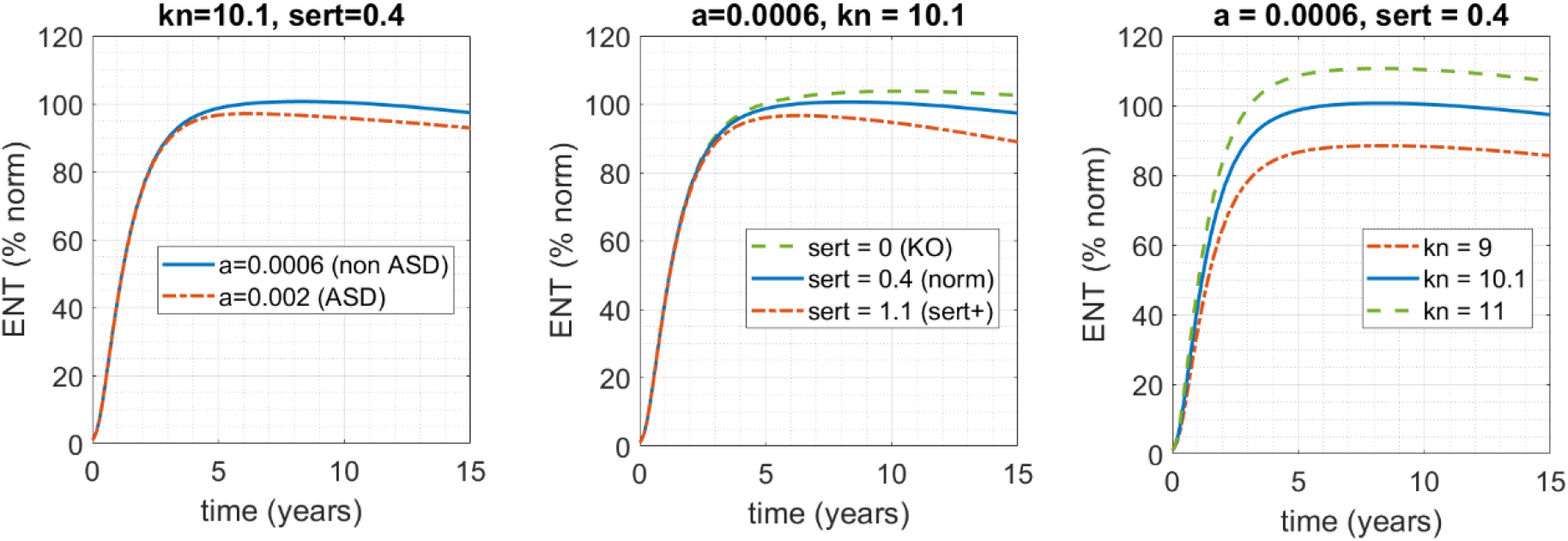
Model-based assessment of the impact of varying the three main factors that functionally connect levels of serotonin in the gut and ENT size. Solid blue line represents the norm (k_n_ = 10.1; *sert* = 0.4; a = 0.0006); dot-dashed red and dashed green lines represent impact of deviation of the norm. (a) The rate of serotonin clearance (*sert* = 0.4) and rate ENT growth as a function of serotonin concentration (*k_n_* =10.1) are fixed; rate of serotonin production over time *a* is varied. (b) Rate of serotonin production over time (parameter *a*) and serotonin-mediated ENT growth (k_n_) are fixed; rate of SERT-mediated serotonin clearance from the gut is varied. (c) Varying rate of serotonin-mediated ENT growth (k_n_), while keeping the other two factors fixed.

As one can see in Figure 4, while all three of these factors can affect the size of ENT by the time t = 5 years, and while each of them can certainly contribute to deviation of ENT size from 100%, neither in this range can solely drive ENT into the 60% of norm range, which would have corresponded to ENT size for the ASD phenotype in the work of Margolis et al. (13). Given the complexity of this condition and its heterogeneity, even for this subset of phenotypes, it is most likely that it is a combination of these factors that would result in the critical phenotype. That is, it is most likely that that there is intrinsic population variability with respect to each of these parameters, and it is a combination of the three rather than a single one that translate into ENT hypoplasia of sufficient severity to manifest in an ASD-like phenotype.

To assess the impact of the combination of these factors, we do a parameter sweep over these 3 properties – *sert* (rate of SERT-mediated serotonin uptake in the gut), *k_n_* (rate of serotonin-mediated ENT growth) and *a* (rate of serotonin production over time in the gut) – and record corresponding size of ENT relative to norm at t = 5 years (Figure 5).

**Figure 5.**
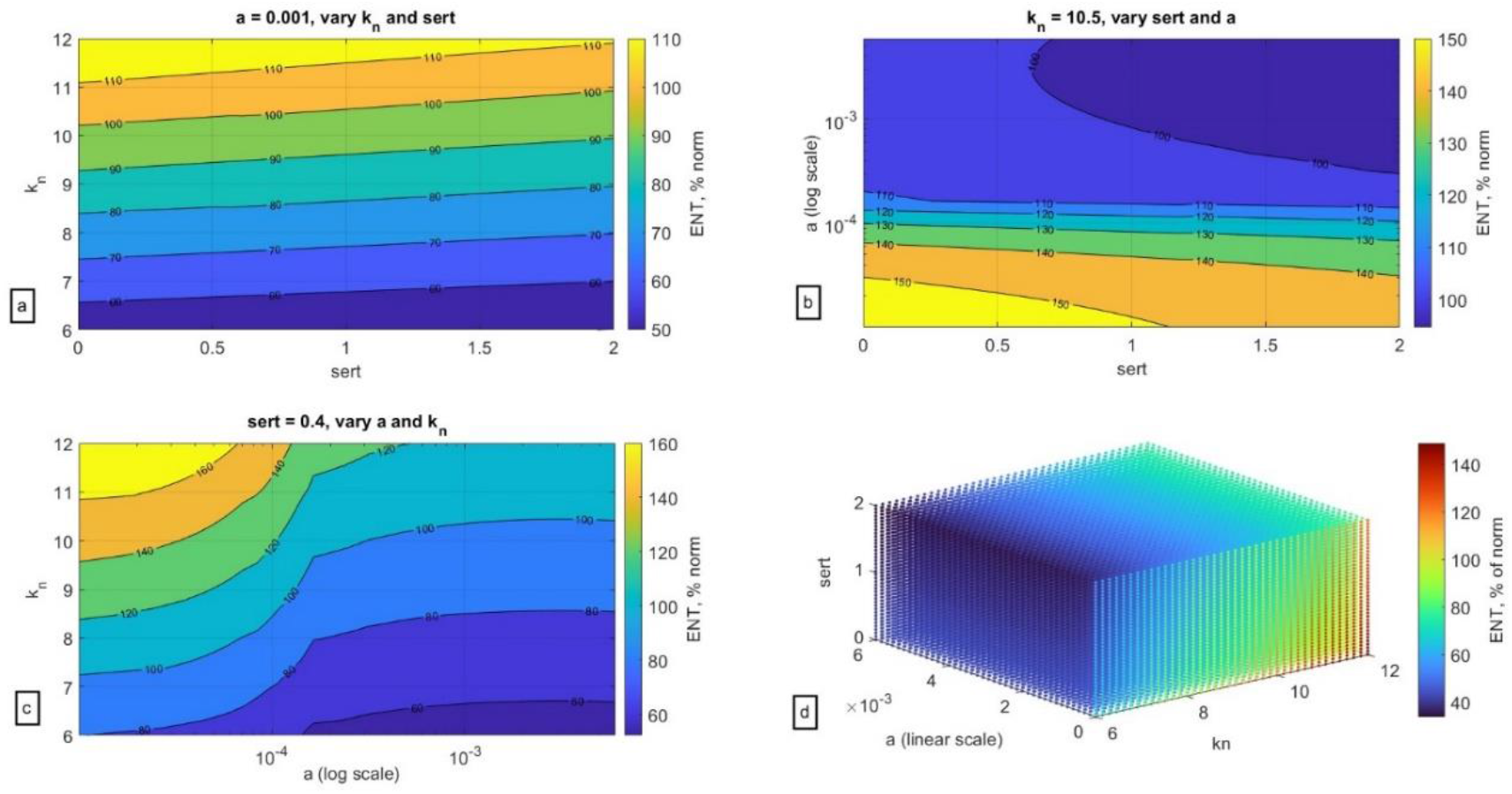
Impact of variation of the three key parameters on ENT size relative to norm (ENT = 100%) at t = 5 years. (a) Holding parameter *a* (serotonin synthesis) constant, (b) holding parameter k_n_ (serotonin-mediated ENT expansion) constant, (c) holding parameter *sert* (SERT-mediated serotonin uptake) constant, (d) three-dimensional plot of variation of all three parameters and their impact on ENT size.

As can be seen in Figure 5, a confluence of factors can indeed significantly affect the size of ENT (here collected at t = 5 years old) more so than each individual factor by itself. As can be seen in Figure 5a, even when serotonin production (parameter *a*) is fixed, low rate of serotonin-mediated ENT growth (parameter k_n_), or high rate of SERT activity (*sert*) can result in significantly lower ENT size. In Figure 5b once can see that when k_n_ is fixed, it is the rate of serotonin production rather than SERT expression that determines ENT size. In Figure 5c, fixing *sert* shows that k_n_ is more important when *a* is high to ensure normal size of ENT but is less impactful for low values of *a*. Finally, in Figure 5d one can see the impact of confluence of all three of these factors, and their relationship with each other as they translate into ENT size. It is possible that, if there is indeed a mechanistic link between ENT size and ASD phenotype, then dynamics reported in Figure 5d may be a reflection of the ASD spectrum, with ENT size relative to the norm potentially reflecting degree of severity of the condition. This hypothesis of course remains to be rigorously evaluated.

## Discussion

This work describes a mathematical model that qualitatively captures results reported by Margolis et al. (13), where the authors showed that at least in Ala56 mice, who overexpress SERT receptor and who exhibit behaviors that broadly align with core diagnostic features of ASD, hypoplastic ENT as a result of increased activity of SERT in the gut is predictive of ASD phenotype. The authors also show that pharmacological interventions that compensate for overexpression of SERT can restore ENT size even in Ala56 mice, and these animals have improvements in behavioral patterns, suggesting that ENT hypoplasia may have a causal link for at least a subset of ASD phenotypes.

Scaling the model from mouse to human suggested that just alterations in SERT expression may be insufficient to result in ENT hypoplasia and instead there exist three key mechanisms that affect this process: rate of serotonin synthesis, rate of serotonin clearance from the gut (such as through SERT receptors), and rate of serotonin-mediated ENT growth. Assuming inter-individual variability with respect to these parameters and conducting a simulation that varies all three of these factors revealed that one can get a “spectrum” of projected ENT size, which perhaps can be correlated to ASD severity, an intriguing hypothesis that requires empirical verification.

More broadly, given the impact of genetics on ASD (3,4,18), within the framework of this analysis, the three key classes of mechanisms identified by the model may reflect a combination of genetic factors, such as serotonin production, clearance and impact on ENT, and it is their confluence that could result in ASD phenotypes that are mechanistically linked to ENT hypoplasia. Interestingly, studies have shown elevated incidence of ASD in children of mothers who took SSRIs during pregnancy, which seems to contradict results reported by (13), who report that in Ala56 mice, administration of SSRIs during pregnancy compensated for excessive activity of SERT and thus mitigated the effects of overexpressed SERT on ENT. It is possible that this effect is a confounder: if the mother has serotonin-mediated depressive symptoms, she may already pass along genetic variant that may pre-dispose the fetus to ASD; it is also possible that there additionally may be an environment where fetal exposure to serotonin during development is decreased (such as has been observed for pre-natal exposure to nutrient limitations, for example (19)), potentially contributing to ENT hypoplasia. If that is the case, then the association is between parental depression, which may be genetic on both sides, that necessitates SSRI administration even during pregnancy, and not the SSRIs themselves, that may be revealed by these observations. While some studies suggest that indeed, the odds of an ASD diagnosis are higher among children whose mothers had a lifetime history of bipolar disorder or depression (20), this hypothesis remains to be evaluated more thoroughly to determine the impact of relevant confounders.

### Potential mechanistic link between ENT hypoplasia and CNS

Of course, an important link that is still unclear is how hypoplastic ENT translates into effects on the CNS. Vagus nerve (VN) is the primary link between ENT and CNS, consisting of afferent VN fibers going “up” from intestinal wall to the brain and accounting for 90% of all VN fibers, and the remaining 10% of efferent VN going “down” from brain to the gut. Afferent VN fibers impact signals relating to inflammation, satiety and energy metabolism, while efferent VN nerves affect secretion of gastric acid and digestive enzymes, as well as gastric capacity (21). Some studies have suggested that vagus nerve stimulation (VNS) may be a promising adjuvant therapy for ASD and other neurodevelopmental disorders (22).

Additionally, mitochondrial dysfunction and impaired energy metabolism may be observed in ASD patients (23,24). Given that the brain accounts for nearly a fifth of the body’s metabolic demands, and that neurons depend on oxidative phosphorylation, impairments in mitochondrial function would make a developing brain uniquely susceptible to neurodegenerative disorders (24). Interestingly, it appears that children prenatally exposed to valproic acid (VPA), a medication used for epilepsy and mood swings, might have an increased likelihood of being diagnosed with autism (25–27), and that VPA may promote mitochondrial dysfunction at least in primary human hepatocytes in vitro (28). Finally, Liu et al. (29) have shown that VPA administration during pregnancy in a rat model additionally affected gut microbiota, with pre-natal exposure to VPA mimicking not only anatomical and behavioral aspects of an autistic brain but also its microbiota.

Gut microbiota has additionally been a focus of intense study in the ASD field. Desbonnet et al. (30) reported that germ free mice tend to exhibit atypical social behaviors, including avoidance of social interactions and excessive grooming. Some bacterial pathogens, such as *Clostridia, Bacteriodetes*, and *Desulfovibrio*, produce propionic acid, which interferes with serotonin synthesis in the gut (31); other bacteria species have been reported to differ in gut microbiota between autistic and non-autistic children as well (32). Impaired metabolism of tryptophan, a precursor to serotonin, has also been reported to be associated with an autistic phenotype (33–35).

Together, these observations suggest that maybe the link between ENT and CNS lies through the ability of the vagus nerve to support energy and mitochondrial metabolism in the brain, and a hypoplastic ENT, which occurs from a combination of factors affecting serotonin’s function as growth factor in ENT, may provide insufficient signaling and thereby affect neurological function. These considerations are summarized in Figure 6.

**Figure 6.**
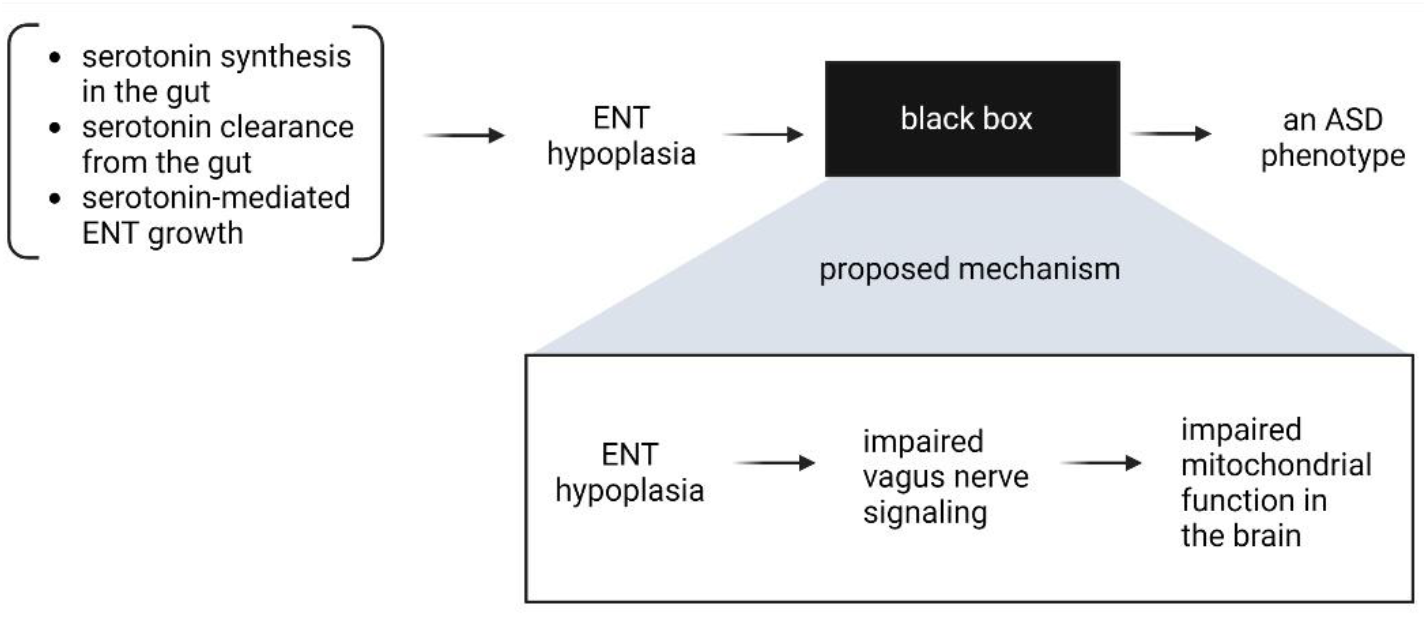
Potential mechanistic connection between ENT hypoplasia and ASD.

Interestingly, transcranial photobiomodulation has been investigated as a possible therapy for ASD (36,37), which is hypothesized to work through improving mitochondrial function in the brain (38,39); studies evaluating this hypothesis are ongoing (40).

The analysis presented here of course has limitations, including numerous assumptions of mechanism translatability between mice and humans with respect to impact of serotonin on ENT. Nevertheless, it does highlight the multi-factorial nature of ASD even within the framework of a simple two-dimensional mathematical model and provides a possible interpretation of ASD spectrum as a function of ENT size relative to norm. Further investigation of this hypothesis might involve development of a genetic panel that focuses on possible genes that contribute to the three mechanisms described here, and then conducting a prospective trial to evaluate the predictive capacity of such a panel. A complementary analysis should be conducted to further elucidate connection between ENT size and its potential impact on metabolism and mitochondrial function in the brain. While genetic pre-disposition to ASD may not be amenable to change, early interventions aimed to mitigate and compensate potential metabolic deficits in combination to already established behavioral therapy may improve quality of life both for children with ASD and their families.

## Acknowledgements

The author reports no external sources of funding. IK is an employee of EMT Serono, US business of Merck KGaA. Views expressed in this work are the author’s personal views and do not necessarily represent the views of EMD Serono.

## Appendix A.

To capture dynamics of serotonin synthesis capacity over time as described by (Chugani et al. 1999), we propose a fully phenomenological function to capture these data where the constant serotonin synthesis rate in Equation (A1) is replaced by the following expression:

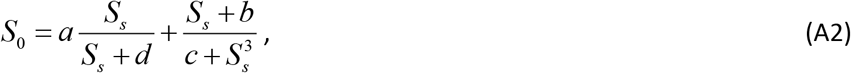

where *S_s_* is systemic serotonin that increases over time. Varying parameter *a* is sufficient to reproduce dynamics reported in Figures 3-5 of the main text.

**Table A1.** Parameters needed to describe serotonin synthesis capacity data for ASD and non-ASD children in the data reported by (Chugani et al. 1999).

| Parameter | Mouse value | Human value |
| --- | --- | --- |
| **a** | .02 | 0.002 for ASD, 0.0006 for non-ASD |
| **b** | 0.01 | 0.01 |
| **c** | 20 | $a/3e-6$ |
| **d** | 1 | 1 |

## Appendix B

Adding qualitative drug dynamics for SSRI fluoxetine, which compensates for hyperactivity of SERT receptor, and prucalopride P(t), which increases serotonin production. A phenomenological depiction of how these mechanisms can be implemented in the proposed model are given below:

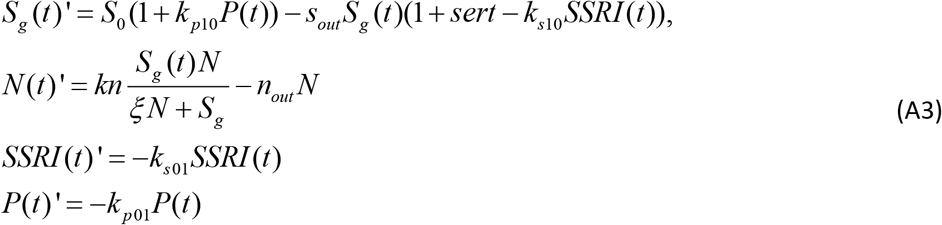

**Table A2.** Additional parameters to describe qualitative drug dynamics and its impact on ENT as reported in (Margolis et al. 2016).

| Parameter | Description | Sample value |
| --- | --- | --- |
| **SSRI fluoxetine** |  |  |
| **k<sub>s01</sub>** | Drug absorption rate into the site of action | 0.1 |
| **k<sub>s10</sub>** | Drug impact on serotonin clearance | 0.12 |
| **Prucalopride** |  |  |
| **k<sub>p01</sub>** | Drug absorption rate into the site of action | 0.1 |
| **k<sub>p10</sub>** | Drug impact on serotonin production | 0.05 |

